# Cell cycling throttles epithelial–mesenchymal transition

**DOI:** 10.64898/2026.11.05.764762

**Authors:** MeiLu McDermott, Anupam Dey, Riddhee Mehta, Adam L. MacLean

## Abstract

Cell cycle phase is an understudied source of variation in gene expression. EMT is a case in point: this cell state transition that drives metastasis has been studied widely across tissue, stimulus, and microenvironmental conditions, yet cell cycle effects are routinely regressed out of single-cell EMT analyses. Here, we compare EMT between G1 and cycling (S/G2/M) cell states in two independent scRNA-seq datasets of TGF-*β*-stimulated MCF10A mammary epithelial cells (varying dose and timepoint). G1 cells consistently show higher expression of EMT genes and upregulated RNA velocities than cycling cells, with the strongest upregulation in G1 EMT intermediate states. Consistent with the go-or-grow hypothesis, EMT and cell cycle programs inversely correlate; moreover, this anticorrelation extends to RNA velocity and strengthens with TGF-*β* dose. We identify a 77-gene G1-specific EMT signature enriched for actin cytoskeletal and extracellular matrix genes that would otherwise be lost by regressing out cell cycle effects. This gene set prompts the hypothesis that, unconstrained by the cytoskeletal demands of mitosis, G1 cells have greater capacity to execute EMT. Further, we developed an EMT gene regulatory model coupled with a cell cycle module, and found through simulation that the model recapitulated this inverse EMT–cell cycle relationship. Using this model, we predict that cycle-selective chemotherapy enriches for the G1 intermediate state, which has high metastatic potential. Cell cycle phase thus ought not be regressed out of EMT states, given its influence on EMT.

## 1 INTRODUCTION

Epithelial–mesenchymal transition (EMT) describes the process by which epithelial cells lose apical-basal polarity and cell-cell adhesion while acquiring the motility and invasiveness of mesenchymal cells (Thiery et al., 2009). Arising in embryonic development and tissue morpho-genesis, EMT is reactivated during cancer progression, where it confers metastatic capacity and contributes to treatment resistance (Dongre & Weinberg, 2019). EMT is not a binary switch. Cells transitioning between epithelial and mesenchymal states also reach stable intermediate states (MacLean et al., 2018; Yang et al., 2020; J. Zhang et al., 2014), the number and identity of which are context dependent (Aiello & Kang, 2019; McDermott et al., 2025). Mathematical models have predicted (and experiments confirmed) the existence of EMT intermediate states, often arising from a core gene regulatory circuit linking TGF-*β* signaling to *SNAIL*, *ZEB*, and the miR-34/miR-200 families (Dey & MacLean, 2025; Lu et al., 2013; Tian et al., 2013). Coupled saddle-node bifurcations in these models generate tri-or tetra-stability (Hong et al., 2015). EMT states are functionally distinct: intermediate rather than fully mesenchymal cells show the greatest plasticity, invasiveness, and metastatic potential in squamous cell carcinoma and pancreatic cancer (Pastushenko et al., 2018; Simeonov et al., 2021), and breast cancer circulating tumor cells are enriched for them relative to the primary tumor (Yu et al., 2013).

Substantial heterogeneity has frustrated efforts to define a common set of genes characterizing EMT across cancer types. Core transcriptional regulators recur across systems — particularly *SNAI1/2*, *ZEB1/2*, and *TWIST1/2* (Lamouille et al., 2014) — yet their expression is context-dependent, and a small number of molecular markers is insufficient to define EMT status (Yang et al., 2020). Downstream of these regulators, the genes identified diverge substantially across studies (Cook & Vanderhyden, 2022), shaped by tissue of origin, EMT-inducing stimulus, and the composition of the local tumor microenvironment (Cook & Vanderhyden, 2020; Tagliazucchi et al., 2023). Single-cell profiling of head and neck tumors, for example, resolved a partial-EMT program built largely from extracellular matrix genes and containing few canonical EMT transcription factors (Puram et al., 2017). This divergence constrains biomarker development and therapeutic targeting: a signature validated in one context may not transfer to another.

One underexplored source of this divergence is cell cycle state (Dong & Blanpain, 2026; Xing & Tian, 2019). An inverse relationship between proliferation and migration has long been documented across cancer types and formalized as the “Go-or-Grow” hypothesis (Akhmetkaliyev et al., 2023), and molecular evidence supports the tradeoff: core EMT transcription factors restrain the cell cycle, with *SNAI1* blocking G1 progression (Vega et al., 2004) and *ZEB2* (*SIP1*) directly repressing cyclin D1 to attenuate the G1–S transition (Mejlvang et al., 2007). EMT inducer TGF-*β* is also documented as an inhibitor of cell cycle (Mukherjee et al., 2010; S. J. Park et al., 2016; Siegel & Massagué, 2003). Although the cell cycle and EMT have, separately, been extensively modeled, EMT models have largely focused on core regulatory circuits without explicit cell cycle dynamics; one Boolean network model, which integrated both, predicted a conflict between mitosis and intermediate EMT stability (Sullivan et al., 2023). Recent work ties this proliferation–migration tradeoff to metastatic outcome. TGF-*β* stimulation of oral cancer cells increased migration in G1 relative to cycling cells (Takahashi et al., 2022), and *in vivo*, elevated *PRRX1* promoted invasion while repressing the cell cycle, with the most metastatic cells occupying an intermediate state at intermediate *PRRX1* levels (Jiménez-Castaño et al., 2026). This coupling is not confined to cancer: in renal fibrosis, a partial EMT program induced cell cycle arrest (Lovisa et al., 2015). In MCF10A cells specifically, Hu et al. (2026) recently reported that EMT couples to cell cycle phase, identifying separable transitions in G1 and G2/M.

Despite the complex relationship between EMT and cell cycle, cell cycle signal is nonetheless routinely treated as a confound in single-cell EMT studies, including those generating new EMT scRNA-seq datasets (Cook & Vanderhyden, 2020; S. Kim et al., 2025; Panchy et al., 2022; Pastushenko et al., 2018; Paul et al., 2023; Simeonov et al., 2021; Wang et al., 2023; Winkler et al., 2024), with phase scores regressed from the expression matrix or cell cycle genes excluded before dimensionality reduction. An axis of interest is thereby removed before analysis begins. Where the EMT–cell cycle relationship has been examined, it has been characterized as a static association between phase and EMT state, leaving open whether cells in different phases traverse the transition at different rates.

Here, we address the interplay between EMT and cell cycle by directly comparing EMT progression in G1 and actively cycling (S/G2/M) cells, using two independent scRNA-seq datasets of TGF-*β*-stimulated MCF10A mammary epithelial cells: one spanning a range of doses, the other a time course. Combining gene expression with RNA velocity, we find that G1 cells reach both higher EMT signature expression and higher EMT velocity than cycling cells, with G1 intermediate-state cells having the highest EMT velocity. We identify a 77-gene G1-specific EMT signature enriched for actin cytoskeleton and extracellular matrix genes rather than canonical EMT regulators, suggesting that G1 cells have greater cytoskeletal bandwidth — with machinery not committed to division — to execute this phenotypic change. Because this signature is phase-specific, the EMT genes identified from a population may depend on its cell cycle composition, contributing to the divergence of EMT signatures across studies. Finally, we extend a core EMT model with a bistable cell cycle module and use it to predict how TGF-*β* exposure shapes the population-level response to cycle-selective chemotherapy.

## 2 METHODS

### 2.1 Data sources

Two publicly available single-cell RNA sequencing (scRNA-seq) datasets of MCF10A cells stimulated with TGF-*β* were analyzed: one across a range of doses (Panchy et al., 2022; GSE213753) and one across a range of timepoints (Paul et al., 2023; GSE194019), hereafter the dose and timepoint datasets.

### 2.2 Sequence alignment

Both datasets were re-aligned to obtain spliced and unspliced read counts for RNA velocity analysis. A *splici* (spliced + intron) index was constructed from the GRCh38 human reference genome using salmon (Srivastava et al., 2019).

For the dose dataset, raw reads (SRX17639978) were downloaded from the NCBI Sequence Read Archive (SRA) (Leinonen et al., 2011) and converted to FASTQ with fasterq-dump from the SRA Toolkit. Reads were mapped to the *splici* index and quantified with alevin-fry (He et al., 2022). Cell barcodes were filtered against the 10x Genomics Single Cell 3’ v2 whitelist, and experimental barcodes for each cell were obtained from the original publication’s supplementary materials.

For the timepoint dataset, raw reads (SRX14892147) were downloaded and converted to FASTQ as above. Because these data were generated with Illumina SureCell 3’ chemistry, FASTQ files were reformatted for alevin-fry following a published vignette (Tekath & Dugas, 2021) and corrected with umi_tools (Smith et al., 2017), allowing up to one error per barcode and an edit distance of 1 per linker segment. Reads were then mapped and quantified as above. Each timepoint was sequenced as a separate library and was therefore processed separately before merging.

### 2.3 scRNA-seq data preprocessing and normalization

All data were processed with scanpy (Wolf et al., 2018). Because the datasets were generated with different sequencing platforms, cells with outlying total counts, gene counts, or mitochondrial percentage were removed using dataset-specific thresholds (Supp. Table 1), and doublets were removed with Scrublet (Wolock et al., 2019). Counts were normalized to 10,000 per cell and log(*x* + 1)-transformed. Cell cycle phase was scored with scanpy’s score_genes_cell_cycle using the gene sets of Tirosh et al. (2016), following the Seurat approach (Satija et al., 2015). Total counts were not regressed out, as cell size correlates with cell cycle phase. Mitochondrial percentage was regressed out for the dose dataset but not for the timepoint dataset, in which it was strongly correlated with timepoint. Highly variable genes were then selected for downstream analysis, and expression was scaled to unit variance.

### 2.4 Cell clustering and EMT score

Principal component analysis (PCA) and nearest-neighbor graph construction were performed with scanpy defaults (50 components, 15 neighbors), and cells were clustered with the Leiden algorithm (Traag et al., 2019) at dataset-specific resolutions (Supp. Table 1). Differentially expressed (DE) genes were identified for each cluster versus all other cells by Wilcoxon rank-sum test with Benjamini–Hochberg correction (adjusted *p <* 0.01; maximum 500 genes per cluster). Cells were visualized with UMAP (McInnes et al., 2020).

EMT gene expression was quantified per cell as an EMT score, computed with the rank-based method UCell (Andreatta & Carmona, 2021) on all filtered genes using the Hallmark EMT gene set from the Molecular Signatures Database (MSigDB) (Liberzon et al., 2015; Subramanian et al., 2005). Clusters were labeled using cell cycle phase, canonical EMT marker expression, and EMT score.

### 2.5 NMF

Non-negative matrix factorization (NMF) was performed on filtered, total-count-normalized counts with scikit-learn’s NMF (Pedregosa et al., 2011). The number of components was increased until an additional component no longer corresponded to an EMT or cell cycle state, and components were annotated by their top-ranked genes. This yielded five components for the dose dataset (epithelial, intermediate, mesenchymal, G1, and cycling) and four for the timepoint dataset (epithelial, intermediate, mesenchymal, and cycling), in which G1 cells were captured by the intermediate component.

### 2.6 RNA velocity with Dynamo and CellRank

Genes were retained for velocity analysis if they had at least 50 combined spliced and unspliced counts and were detected in at least 30 cells in each of the spliced and unspliced layers. RNA velocity was then inferred with Dynamo (Qiu et al., 2022) using default parameters.

Two velocity-based gene sets were derived. First, differential velocity genes were identified with scVelo’s rank_velocity_genes (Bergen et al., 2020), which compares velocities in each cluster against all other cells using Welch’s *t*-test with overestimated variance; genes with *t >* 5 were retained. Second, fate-biased genes were identified with CellRank (Lange et al., 2022) using a combined velocity kernel (0.8 velocity, 0.2 expression). Macrostates were computed with the GPCCA estimator, with the number of macrostates set to the number of clusters in each dataset. Fate probabilities toward either the G1 or cycling terminal states were computed with compute_fate_probabilities, and genes correlated with each fate (*q <* 0.01) were identified with compute_lineage_drivers.

### 2.7 EMT score–cell-cycling score space

A cell-cycling score was computed with Scanpy’s score_genes using the S and G2/M genes from Tirosh et al. (2016). Velocities were projected into the two-dimensional EMT score–cell-cycling score space with Dynamo and visualized with streamline_plot. Spearman’s correlation between EMT score and cell-cycling score velocities was calculated across doses and timepoints. Kernel density estimates (KDEs) were computed per cluster with SciPy’s gaussian_kde; the KDE mode of EMT score and cell-cycling score defined each cluster’s center, and the KDE mode of the projected EMT score and cell-cycling score velocities defined the cluster’s vector.

### 2.8 Cell state density and low-density dynamic genes with Mellon and Palantir

Low-density dynamic genes were identified following the Mellon workflow (Otto et al., 2024). Cells were first split into G1 and cycling subsets by phase, and cell-state density was estimated within each subset using Mellon, with the number of diffusion components selected by the elbow method (Supp. Table 1). Trajectories were inferred using Palantir (Setty et al., 2019), with initial and terminal states assigned from cluster identities via early_cell and find_terminal_states. Expression was smoothed with MAGIC (van Dijk et al., 2018), and genes variable in low-density regions were identified from the smoothed expression and Mellon density estimates with Palantir’s run_local_variability. This procedure was applied to four trajectories (G1 and cycling subsets of each dataset) with transitions considered between EMT clusters. For each transition, genes scoring above the elbow of the variability score distribution were retained.

### 2.9 Multi-analysis gene aggregation

G1 and cycling EMT gene sets were defined separately, both using the following procedure. Genes were first required to be differentially expressed in the intermediate and/or mesenchymal state in both the dose and timepoint datasets. These genes were then cross-referenced against two types of analyses, RNA velocity (scVelo differential velocity genes and CellRank fate-biased velocity genes) and low-density transition genes (Mellon with Palantir), and were retained if genes were significant in at least two of the six dynamical analyses (scVelo, CellRank, and Mellon in each dataset). Velocity fits for the resulting genes were manually inspected, and genes with poor fits were excluded.

Gene ontology (GO) analysis of the G1 EMT gene list was performed with DAVID (Sherman et al., 2022), with the background restricted to genes expressed in both datasets.

### 2.10 A mathematical model coupling EMT and cell cycle circuits

The EMT–cell cycle coupled circuit was modeled as a system of differential equations comprising two modules. The EMT module, adapted from the tristable model of Lu et al. (2013), consists of three nodes: miR-200 (R; Eq. 1), the transcription factors ZEB1/2 (Z; Eq. 2), and the input signal TGF-*β* (S). The cell cycle module (C; Eq. 3) represents net expression of pro-cell-cycle progression factors. In brief: R is inhibited by S and Z; Z is activated by S, inhibited by R, and self-activating; and C is self-activating and inhibited by S and Z. Inhibitions of the cell cycle from TGF-*β* and ZEB1/2 are supported by literature (Hugo et al., 2013; Mejlvang et al., 2007; S. J. Park et al., 2016). Parameter values were derived using the method of Dey and Barik (2021) and are described in Supp. Table 2. The model equations are:

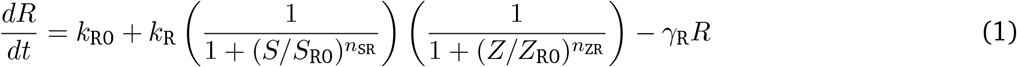

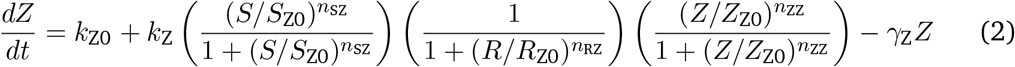

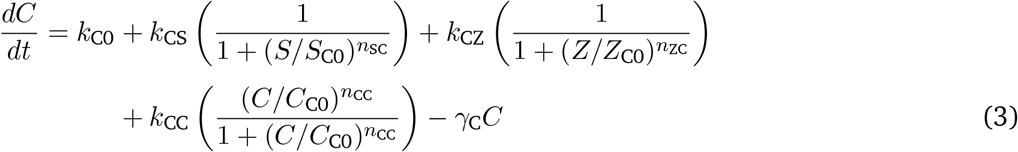

In the EMT module, regulatory inputs are combined multiplicatively (AND logic), such that high expression of a node requires all of its inputs to be permissive (high activator and low inhibitor levels); this is consistent with prior core EMT networks (Lu et al., 2013). In the cell cycle module, inputs are combined additively (OR logic), such that high expression of C can be achieved through any single permissive input. This difference reflects node composition: inputs to R and Z act on a single species, whereas inputs to C act on an aggregate of genes and can therefore contribute independently.

Bifurcation diagrams were constructed in MATLAB by numerically solving the steady-state equations. For each value of the signal *S*, fixed points were located by a grid-based sign-change search and then refined, and their stability was determined from the eigenvalues of the Jacobian. Nullclines were computed as the zero-level curves of each differential equation evaluated on a two-dimensional grid of the state variables, with the independently computed fixed points at their intersections.

### 2.11 Stochastic simulations of the EMT–cell cycle model

Single-cell trajectories were simulated by extending the ODE model to a system of stochastic differential equations (SDEs) with additive noise:

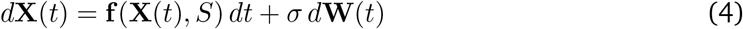

where **X**(*t*) = [*R, Z, C*]*^T^*is the state vector, **f** (**X**(*t*)*, S*) is the deterministic drift given by Eqs. 1–3, *σ* is a constant noise strength, and *d***W**(*t*) is a three-dimensional Wiener process with independent increments *d***W**(*t*) *∼ N* (**0***, dt* **I**).

For each value of *S* (TGF-*β* dose), 1,000 cells were simulated with the Euler–Maruyama scheme using a fixed time step Δ*t* = 0.001 and total simulation time *T* = 1000. At each step, each state variable *X ∈ {R, Z, C}* was updated as

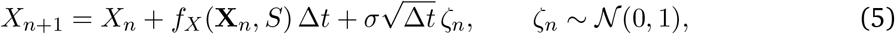

with *σ* = 10 and *ζ_n_* drawn independently for each variable and cell.

At selected timepoints, each cell was assigned to the nearest stable steady state of the deterministic model at the corresponding *S*, with unstable steady states as boundaries. This classified cells by *Z* (epithelial, intermediate, or mesenchymal) and *C* (G1 or cycling). Aggregating these assignments across cells yielded population state distributions over time.

### 2.12 Stochastic simulations with cycle-selective chemotherapy

Cycle-selective chemotherapy was modeled as an instantaneous, state-dependent death event at *t* = 100. Cells classified as cycling at that time were killed independently with probability 0.75, whereas G1 cells were unaffected. Dead cells were removed from the population and excluded from all subsequent integration and state distributions, and were not replaced. Simulations were otherwise identical to those described above.

## 3 RESULTS

### 3.1 G1 cells reach higher EMT scores than cycling cells

To investigate the relationship between EMT and cell cycle states, we analyzed two datasets in which MCF10A mammary epithelial cells were stimulated with TGF-*β* over a range of doses (Panchy et al., 2022) or timepoints (Paul et al., 2023). Both datasets contain cells in states spanning the EMT spectrum and in G1, S, and G2/M phases of the cell cycle (hereafter, S and G2/M phases are together termed “cycling”). Single cells were clustered by transcriptional profile to identify cell states (Fig. 1a). Each cell was assigned a cell cycle phase and then scored for EMT expression using the MSigDB Hallmark EMT gene set (Liberzon et al., 2015) to give EMT scores (Fig. 1b and Methods). Cell states were readily identifiable and thus labeled by their predominant cell cycle phase and EMT state. In both datasets, EMT scores were significantly higher for G1 cells than cycling cells, a difference evident in both the intermediate and mesenchymal states (Fig. 1b and Supp. Fig. 1).

**Figure 1:**
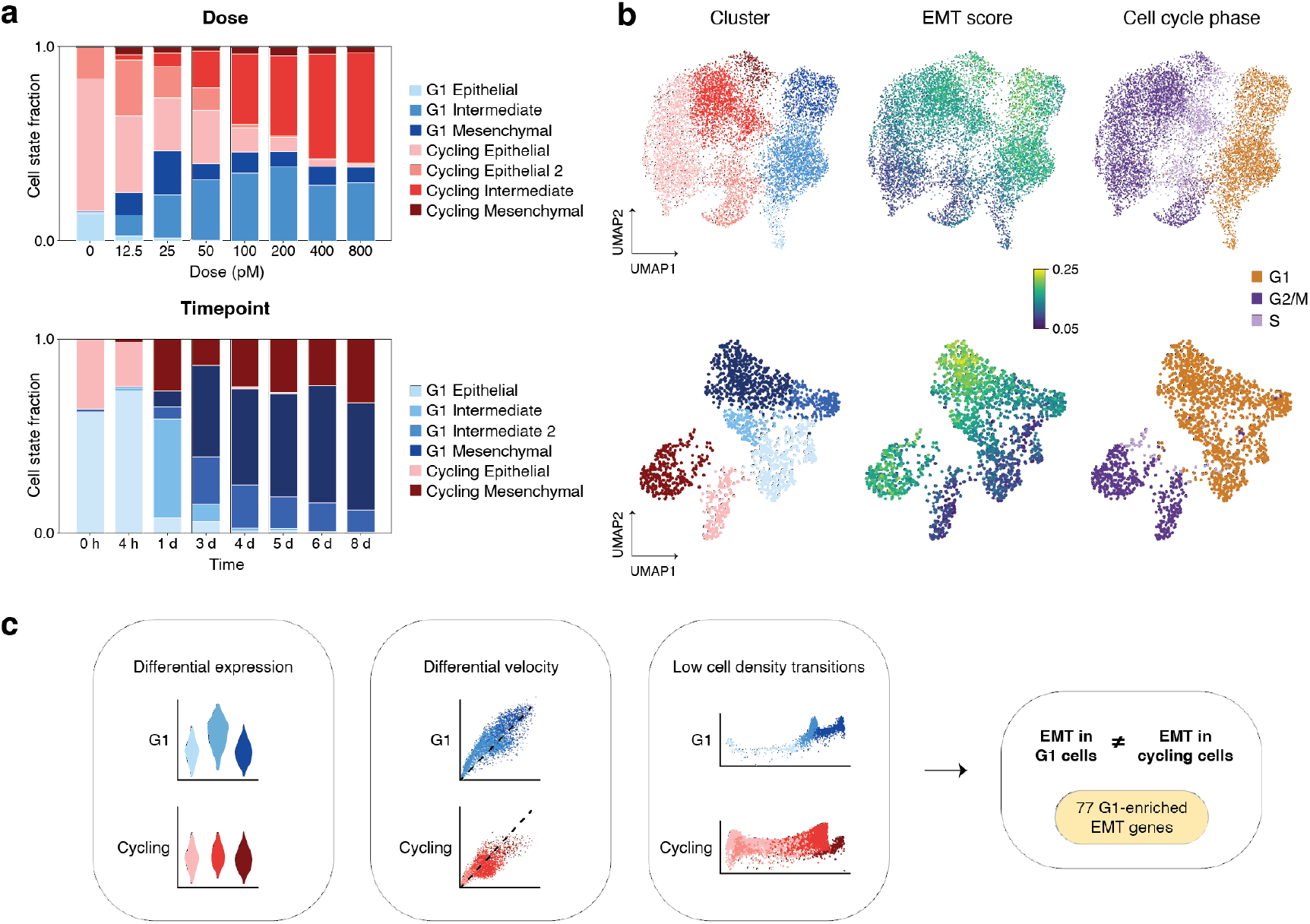
EMT trajectories in MCF10A cells stimulated with TGF-β at varying doses and timepoints. **a)** Cluster proportions across TGF-β dose and time conditions.**b)** UMAP embeddings for each dataset, colored by cluster, EMT score, and cell cycle phase. G1 cells attain the highest EMT scores. **c)** Pipeline combining differential expression, differential RNA velocity, and low-density transition analyses to identify 77 genes associated with G1-specific EMT.

Cluster patterns and cell proportions at dose/timepoint 0 differed between the two datasets, reflecting divergence between the untreated epithelial cell states (Supp. Fig. 2). Many genes were differentially expressed between the two datasets under these control conditions, even after normalization, batch correction, and filtering for only genes expressed in both datasets. This divergence may arise from cell-line drift (the two studies sourced MCF10A cells from different providers) and different culture conditions, including EGF, insulin, and hydrocortisone levels (Supp. Table 3). The timepoint dataset also included a 16-hour serum-free step; the resulting depletion of growth factors is a plausible driver of its lower cycling-to-non-cycling ratio in the untreated condition (Fig. 1a). These compositional differences complicate direct integration of the two datasets, but strengthen the observation that EMT scores are elevated in G1 cells in both datasets.

**Figure 2:**
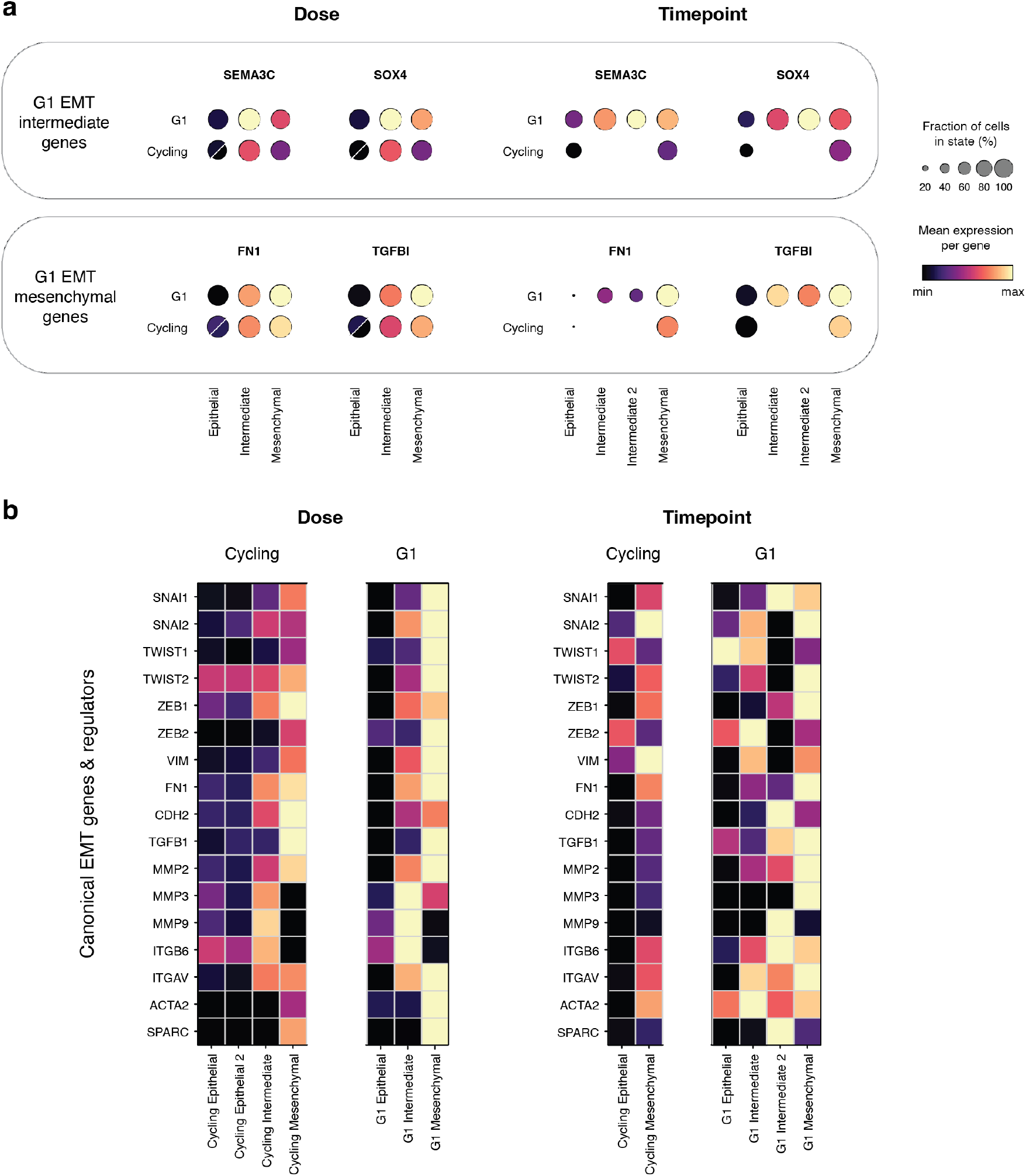
Canonical EMT genes show higher expression in G1 cells than in cycling cells. **a)** Representative genes for EMT intermediate states (SEMA3C, SOX4) and mesenchymal states (FN1, TGFBI) peak in expression in G1 cell states. Split circles represent the two epithelial clusters in the dose dataset. **b)** Gene expression shown by state for a curated panel of 17 canonical EMT markers and regulators.

To identify the transcriptional basis of this difference in EMT scores, we combined three analyses: differential expression analysis (Wolf et al., 2018), to find genes with distinct expression between G1 and cycling cells; differential RNA velocity analysis with scVelo (Bergen et al., 2020) and CellRank (Lange et al., 2022), to find genes with distinct regulatory dynamics; and low-density transition analysis with Mellon (Otto et al., 2024), to find genes marking distinct transition dynamics (Fig. 1c). Together, these analyses defined a 77-gene G1 signature, distinguishing G1 from cycling cells during EMT.

### 3.2 Canonical EMT genes peak in G1 cells

Differential expression between G1 and cycling cells revealed multiple canonical EMT genes consistently elevated in the G1 intermediate and mesenchymal states across both datasets (Fig. 2a). Representative examples include *SEMA3C* (semaphorin 3C) and *SOX4* (SRY-box transcription factor 4) in the intermediate states, and *FN1* (fibronectin 1) and *TGFBI* (transforming growth factor beta-induced) in the mesenchymal states. More broadly, a curated panel of 17 canonical EMT markers and regulators (Lamouille et al., 2014) showed consistently higher expression in the G1 intermediate and mesenchymal states than in the equivalent cycling cell states (Fig. 2b).

Although the overall pattern of G1 EMT upregulation is present in both datasets, expression is more complex in the timepoint dataset: known EMT genes are not uniformly upregulated in intermediate and mesenchymal states. This may reflect the dataset’s finer EMT-state structure (two intermediate states) and denser temporal sampling, which better captures transient expression as cells transition through EMT.

As an additional approach to gain insight into these transcriptional differences, we performed non-negative matrix factorization (NMF) on each dataset, identifying five components in the dose dataset and four in the timepoint dataset (Supp. Fig. 3). In both datasets, individual components corresponded to EMT states (epithelial, intermediate, mesenchymal) and cell cycle phase. Components associated with the intermediate and mesenchymal states were more enriched in G1 than cycling cells, thus corroborating the differential expression results for canonical EMT genes.

**Figure 3:**
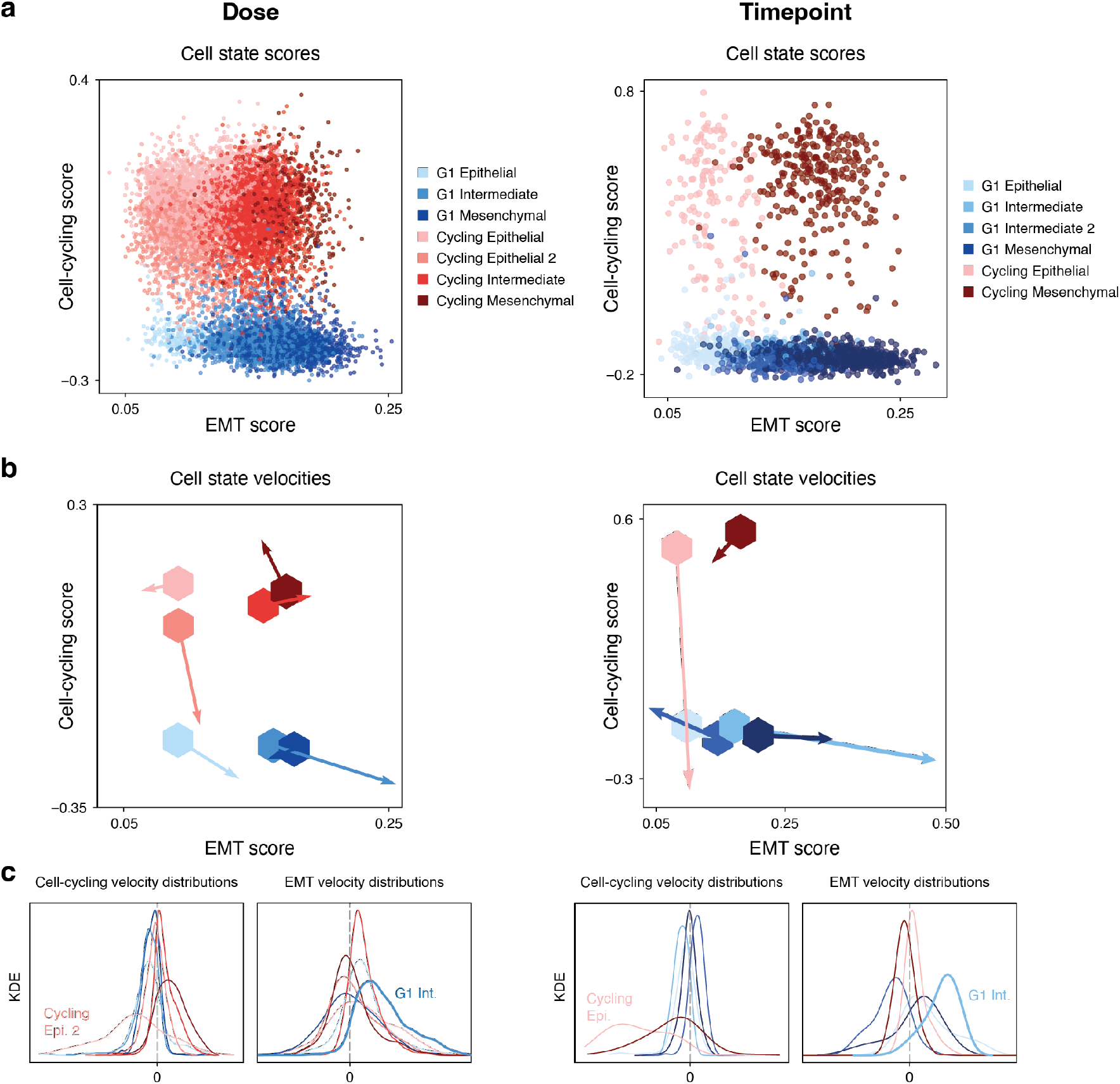
a) For each dataset, EMT scores and cell-cycling scores are plotted per-cell and colored by cluster. **b)** Cluster centers (hexagons) are shown on the same EMT score—cell-cycling score axes, with arrows indicating the overall RNA velocity of each cluster (from the modes of the velocity distributions). **c)** Marginal distributions of the velocity vectors along the EMT score and cell-cycling score axes.

### 3.3 G1 cells upregulate EMT genes more strongly than cycling cells

To decompose the transcriptional effects of progression through the cell cycle versus progression through EMT, we analyzed the two processes jointly, representing each cell in a two-dimensional space defined by two independent scores computed from non-overlapping gene sets: the EMT score and the cell-cycling score. The cell-cycling score is computed per cell from the combined set of canonical S- and G2/M-phase genes (Tirosh et al., 2016). Higher values indicate greater expression of cell cycle genes in that cell. These per-cell scores recapitulate cluster labels (Fig. 3a, Supp. Figs. 4 and 5). We then projected each cell’s RNA velocity into this space (Supp. Fig. 6), giving each cell a position and direction of movement in the joint EMT–cell-cycle plane.

**Figure 4:**
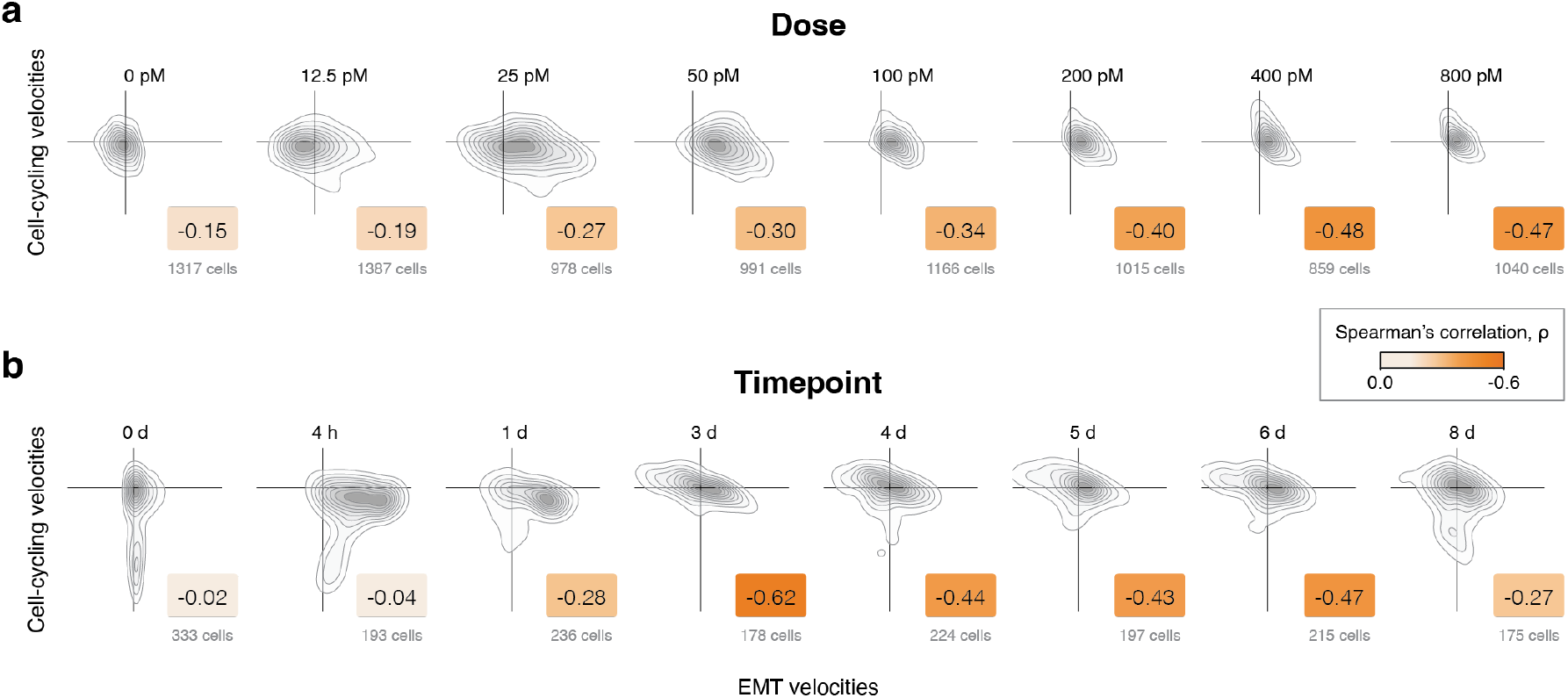
RNA velocities plotted across doses or timepoints on the EMT score—cell-cycling score axes. Negative correlations are quantified by Spearman’s ρ. **a)** Dose dataset: the anticorrelation strengthens with dose, plateauing at the highest doses. **b)** Timepoint dataset: the anticorrelation peaks at middle timepoints. Axis limits are consistent within each dataset.

**Figure 5:**
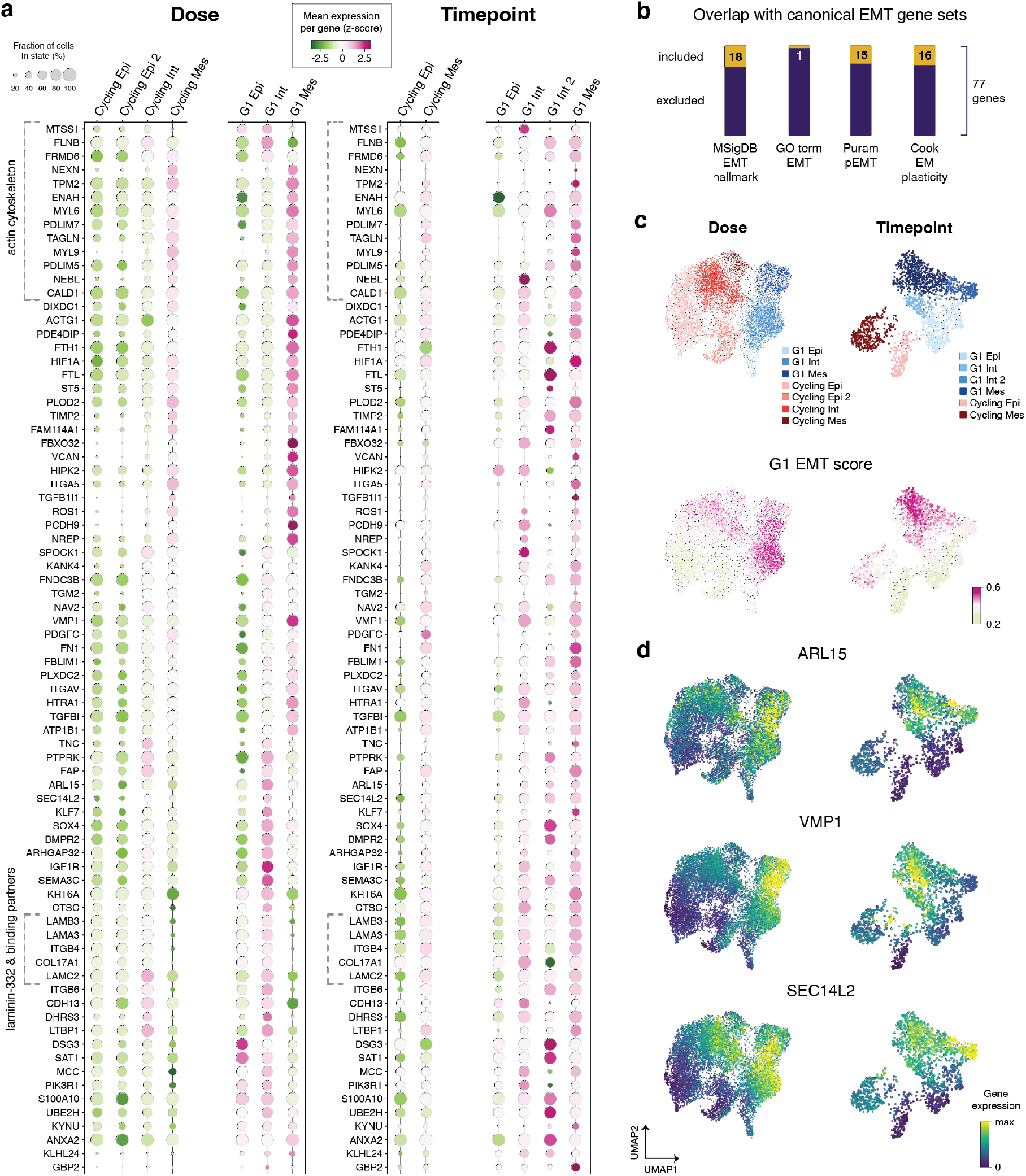
Gene signature for G1-specific EMT. **a)** Expression of the 77-gene signature across dose and timepoint datasets. **b)** Overlap between the G1 EMT gene set and canonical EMT gene sets: a majority of the predicted G1 EMT genes are absent from these lists. **c)** Combined expression of the 77 genes as a G1 EMT score, showing elevation in the G1 intermediate and mesenchymal states. **d)** Expression of three genes absent from canonical EMT gene sets: ARL15, VMP1, and SEC14L2.

**Figure 6:**
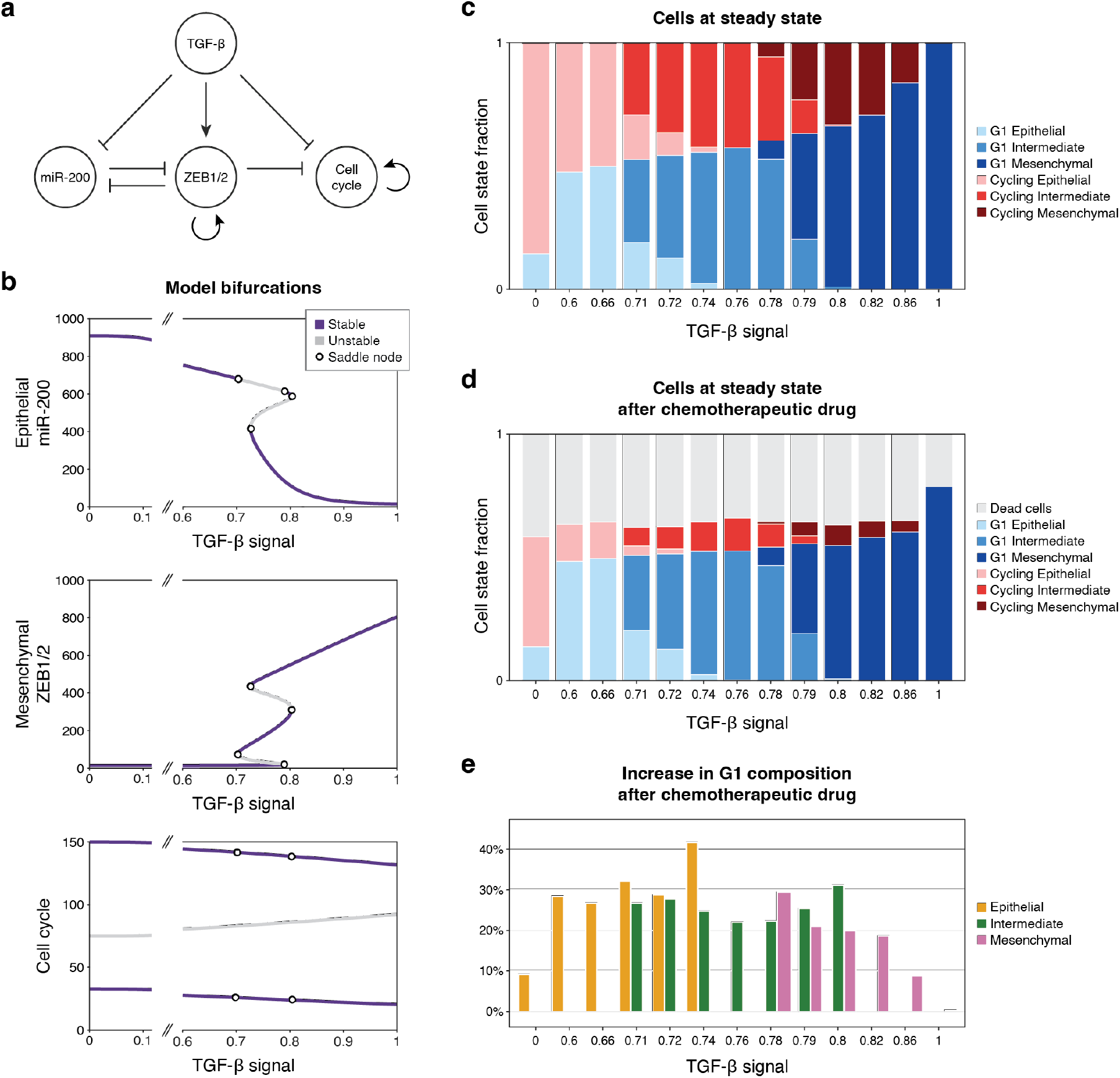
A model of coupled EMT and cell cycle dynamics. **a)** Network topology and regulatory interactions in the model. **b)** Bifurcation diagrams of miR-200, ZEB1/2, and cell cycle across TGF-β signal. For the chosen parameter set, the system is multistable between TGF-β ≈ 0.7 and 0.8, bounded by saddle-node bifurcations; the cell cycle is bistable over the entire TGF-β range. **c)** Steady-state composition resulting from stochastic simulations, initialized with epithelial cells split evenly between G1 and cycling states. **d)** As in (c), with a simulated chemotherapeutic agent killing 75% of cycling cells early in their trajectory. **e)** The resulting increase in total G1 composition relative to the untreated distributions from (c).

For each cluster, we summarized per-cell velocity vectors into a single net vector (Fig. 3b), computed as the mode of their kernel density estimate (Fig. 3c). Individual cells vary in EMT and cell-cycling velocity, but these distributions are unimodal with well-defined peaks, justifying the use of a single summary vector per cluster.

In both datasets, a G1 EMT intermediate state shows the largest EMT velocity (abscissa; Fig. 3b): the strongest EMT upregulation occurs in cells not actively dividing. In these EMT-high intermediate states, net cell-cycling velocities (ordinate) are slightly negative, indicating that cells in these clusters are not likely to be entering the cell cycle. Intriguingly, the two G1 intermediate states in the timepoint dataset (Intermediate and Intermediate 2) diverge in EMT velocity: while the one noted above shows the largest magnitude observed, the other has near-zero EMT velocity, suggesting that it has reached a terminal or relatively more stable (deeper basin of attraction) intermediate state. These results indicate that EMT upregulation happens primarily in cells already in G1, which then continue through the EMT program and are unlikely to enter the cell cycle.

In both datasets, epithelial-state cycling cells show large negative cell-cycling velocities (Fig. 3b), indicating that the majority of epithelial cells actively exit (or fail to enter) the cell cycle upon TGF-*β* stimulation. Mesenchymal-state cycling cells (a smaller population) do not show this negative velocity, consistent with the documented cytostatic effect of TGF-*β* on epithelial cells, including in mammary epithelial models (Donovan & Slingerland, 2000; Y. Zhang et al., 2017).

### 3.4 EMT and cell-cycling velocities are inversely correlated

EMT velocity and cell-cycling velocity are inversely correlated at the single-cell level. Given that these are velocities (changes in transcriptional state rather than transcriptional states themselves), this reflects an active tradeoff between the two programs, reinforcing the conclusion above that EMT upregulation is concentrated in cells that are not actively cycling. On the same EMT–cell-cycling score axes, their joint distributions show negative Spearman correlations in both datasets (Fig. 4): cells upregulating EMT concurrently exit the cell cycle, and vice versa. The strength of this anticorrelation varies with TGF-*β*, with the two datasets giving complementary views. In the dose dataset, the anticorrelation strengthened monotonically with dose, consistent with TGF-*β* acting as both an EMT inducer and a cell-cycle repressor (Mukherjee et al., 2010), until the highest doses, where the correlation held constant. At these highest doses, positive EMT velocity remains, with many cells in intermediate EMT (Fig. 1a), suggesting that some high-dose cells have not yet reached equilibrium. In the timepoint data, the anticorrelation peaked at middle timepoints and weakened at early and late times. Together, the two datasets indicate that the EMT–cell-cycle inverse relationship is strongest while EMT is actively progressing.

RNA velocities of untreated cells differed between datasets, reflecting their distinct baseline compositions (Fig. 4; 0 pM, 0 d). The untreated cells from the dose dataset show a unimodal velocity distribution, whereas the timepoint 0 cells contain a subset of cells with strongly negative cell-cycling velocities, indicating active exit from cycling. This is consistent with the untreated timepoint being predominantly G1 (Fig. 1a), likely reflecting its serum-free preculture in contrast to the majority-cycling dose 0 population. The inverse EMT–cell-cycling velocity relationship nonetheless holds in both datasets, indicating that it is robust to their differing baseline cell-cycle composition.

### 3.5 Gene set characterizing G1 EMT is enriched in actin cytoskeletal remodeling

Given the upregulation of EMT genes in G1 cells, we sought to identify the underlying transcriptional differences by combining three analyses: differential expression between cell states, differential RNA velocity (DV) between G1 and cycling cells, and low-density cell-state transition analysis along the EMT trajectory (example genes in Supp. Fig. 7). We performed DV with two methods that differ in their assumptions: scVelo, which tests each cell state individually (Bergen et al., 2020), and CellRank, which uses fate bias to infer genes associated with progression toward G1 or cycling mesenchymal states (Lange et al., 2022). Because both CellRank and the Mellon low-density analysis (Otto et al., 2024) assume a trajectory, we analyzed G1 and cycling cells separately, supported by G1 cells largely undergoing EMT without entering the cell cycle (Fig. 3b). Mellon relies only on expression, meaning this method can capture transition genes that velocity analyses miss due to low capture. Genes were included in the G1 EMT gene set if they were differentially expressed in an intermediate or mesenchymal state in both datasets and were significant in at least two of the six dynamical analyses (scVelo, CellRank, and Mellon in each dataset; Supp. Tables 4 and 5).

The resulting G1 EMT gene set comprises 77 genes (Fig. 5a), enriched for actin cytoskeletal organization genes (GO:0030036, *p*_adj_ = 0.005; Supp. Table 5). This enrichment suggests a “cytoskeletal bandwidth” model of the EMT–cell-cycle inverse relationship: cycling cells — particularly those entering mitosis, when the actomyosin cortex and contractile ring draw heavily on shared cytoskeletal components (Taubenberger et al., 2020) — may be less able to execute the morphological remodeling of EMT than cells in G1 phase.

The G1 EMT signature is dominated by structural effectors of EMT — genes encoding protein machinery that physically remodels the cell and its extracellular matrix contacts. This is reflected in enrichment for extracellular matrix (ECM) and cell adhesion genes (GO:0031012, *p_adj_* = 2 *×* 10*^−^*^12^ and GO:0007155, *p_adj_* = 4 *×* 10*^−^*^14^ respectively; Supp. Tables 6 and 7).

Together, genes included in these terms account for 37 of the 77 genes. Among them are all three laminin-332 genes (*LAMA3*, *LAMB3*, *LAMC2*) and their hemidesmosomal receptors *ITGB4* and *COL17A1*; laminin-332–ITGB4 is linked to EMT at tumor margins in breast cancer (B. G. Kim et al., 2011), and COL17A1 stabilizes laminin-332 to promote EMT and metastasis in lung cancer (Liu et al., 2018). The G1 EMT gene set overlaps partially with known EMT gene lists (Fig. 5b), particularly among extracellular-space genes. A combined “G1 EMT score” summarizing the 77-gene set (constructed similarly to scores above) confirms elevated expression across the G1 intermediate and mesenchymal states (Fig. 5c).

The G1 EMT gene set also includes genes absent from canonical EMT signatures, i.e. genes with important contributions to EMT but whose signal is attenuated or removed by cell-cycle regression. We highlight three examples (Fig. 5d). The GTPase *ARL15* positively regulates TGF-*β*/SMAD signaling and promotes migration and invasion in breast cancer cells (Shi et al., 2022). The endoplasmic reticulum protein *VMP1* builds and maintains cell junctions, a process central to EMT remodeling, and has been reported to both suppress and promote invasion across cancers (Chen et al., 2026; Li et al., 2026; Renna et al., 2024). Knockdown of the lipid-binding protein *SEC14L2* reduces migration in NSCLC cells (Zhou et al., 2024), though it has no direct link to EMT. Together, these genes point to migration- and invasion-relevant biology beyond canonical EMT regulators.

In contrast, applying the same approach to cycling (S/G2/M) cells undergoing EMT yielded few EMT-specific genes; the list was dominated instead by cell cycle genes (Supp. Table 4). The G1 EMT gene set recovers canonical EMT genes alongside ECM and structural effectors, marking the stronger EMT upregulation reached in G1 as a distinct transcriptional program.

### 3.6 A coupled EMT–cell cycle model recapitulates G1 state enrichment during EMT and predicts EMT-dependent chemotherapeutic responses

To extrapolate beyond the data and study how EMT–cell cycle coupling can respond to perturbations, we developed a simple mathematical model coupling EMT and the cell cycle, incorporating the dual effects of TGF-*β* on EMT and the cell cycle into a previously characterized EMT model (Lu et al., 2013; Tian et al., 2013).

The EMT circuit comprises mutually inhibitory miR-200 (epithelial) and *ZEB1/2* (mesenchymal) nodes and predicts a stable intermediate state. To it we added a cell cycle module whose positive feedback loop yields two stable states, representing active cycling and G1 (Fig. 6a). Both edges linking TGF-*β* and EMT to this module are supported by prior work: TGF-*β* is cytostatic in epithelia (Mukherjee et al., 2010; Siegel & Massagué, 2003), and *ZEB2* (*SIP1*) represses the cell cycle through cyclin D1 (Mejlvang et al., 2007). The coupled system reproduces both features of our data (Fig. 6b): ZEB1/2 is tristable at TGF-*β* levels *∼*0.7–0.8, and cell cycle appears bistable across the full range of TGF-*β* levels considered. The full stability landscape is more complex, however, with 15 fixed points but only 3 unique values in *C^∗^* (5-fold degeneracy; Supp. Fig. 8).

To examine population-level behavior, we simulated cells stochastically (Fig. 6c; 1000 cells per TGF-*β* value). All cells were initialized in the epithelial state (high miR-200, low *ZEB1/2*) and split evenly between cycling and G1, leaving TGF-*β* as the only variable across conditions. At steady state, the fraction of intermediate and mesenchymal cells increased with TGF-*β*, as did the fraction of non-cycling (G1) cells. The model thus recapitulates the inverse relationship between EMT and active cell cycling observed in the data.

We next asked how this coupling between EMT and the cell cycle shapes response to cycle-selective chemotherapy. Under the same simulation conditions, we modeled chemotherapy by removing 75% of cycling cells early after stimulation, reflecting the selective cytotoxicity of many conventional chemotherapeutic agents toward cycling cells (Anand et al., 2022) (Fig. 6d). Cycling populations were depleted as expected, but the resulting shift toward G1 depended on TGF-*β*, reported here as the absolute change in G1 fraction among surviving cells (Fig. 6e). At TGF-*β* = 0, the cycling epithelial population recovers over the post-treatment trajectory (Supp. Fig. 9), a recovery progressively lost with increasing TGF-*β*. The G1 intermediate state showed the most consistent increase across the multistable TGF-*β* range, while the epithelial and mesenchymal states varied more. Due to the high metastatic potential of EMT intermediate cells (Pastushenko et al., 2018; Simeonov et al., 2021), this treatment-induced enrichment of the intermediate state could add considerably to metastatic burden. The G1 shift declined again at higher TGF-*β*, where the untreated population was already predominantly mesenchymal and G1-enriched.

This model shows that a minimal circuit model coupling EMT and cell cycle multistability is sufficient to reproduce the inverse EMT–cell cycling relationship observed in our data. It further predicts that cycle-selective chemotherapy enriches for cells in a G1 EMT intermediate state, a population dangerous for its metastatic potential.

## 4 DISCUSSION

We have shown that EMT is elevated in G1 relative to cycling cells. The 77-gene G1 EMT signature is dominated by structural effectors of the transition — actin cytoskeletal and extracellular matrix genes. We propose that a cytoskeletal bandwidth constraint underlies this asymmetry. Cycling cells, particularly those entering mitosis, commit much of their actin cytoskeleton to the cortical and contractile-ring functions required for division, leaving less available for the stress fibers, lamellipodia, and focal adhesions that mesenchymal morphology demands (Datta et al., 2021; Friedl & Wolf, 2010). G1 cells may therefore have greater capacity to execute the EMT program. Treating EMT and cell cycle as separate axes and inferring joint velocities captures both transitions within the same cell, separating how far a cell has progressed from how fast it is changing. By this measure, G1 cells emerge as the population with the highest EMT velocities and exhibit negligible cell-cycling velocities. Such an orthogonal approach could generalize to other biological processes suspected to be confounded by cell cycle phase.

More broadly, an inverse relationship between proliferation and migration, known as the “Go-or-Grow” model (Akhmetkaliyev et al., 2023), has been documented across multiple cancers (Hoek et al., 2008; Jerby et al., 2012; Oliveira et al., 2017), though not all cancers (Thiessen et al., 2025). Our results are consistent with such a tradeoff operating within EMT and suggest cytoskeletal bandwidth as one constraint that could enforce it. The converse reinforces this interpretation: the inferred cycling–EMT gene set contains few EMT effectors, as expected if actin committed to division is unavailable for mesenchymal remodeling.

Cell cycle phase is a contributing axis of EMT heterogeneity, a point on which our results and recent work converge. Analyzing the same dose dataset with protein-level validation, Hu et al. (2026) reported that EMT couples to the cell cycle, with distinct transitions in G1 and in G2/M; our results complement these, showing that G1 cells upregulate an EMT program and identifying genes associated with this program. Studies in other systems tie the coupling of mesenchymal state and cell cycle exit to dissemination *in vivo*. In oral squamous cell carcinoma, TGF-*β*-treated G1 cells are more migratory than cycling cells *in vitro*, and xenograft lymph node metastases are predominantly G1, whereas primary tumors contain both populations (Takahashi et al., 2022). In murine breast cancer, *ZFP281* maintains early disseminated cells in a dormant, invasive, mesenchymal-like state (Nobre et al., 2022). G1-arrested murine breast cancer cells undergo greater EMT via ribosome biogenesis (Prakash et al., 2019), pointing to a second resource that G1 cells can redirect toward EMT alongside our proposed cytoskeletal bandwidth.

Despite this body of evidence as to the importance of the cell cycle’s biological effects, correcting for cell cycle remains common practice in single-cell RNA sequencing studies (Cook & Vanderhyden, 2020; S. Kim et al., 2025; Pastushenko et al., 2018; Simeonov et al., 2021; Wang et al., 2023; Winkler et al., 2024), including the original analyses of both datasets used here (Panchy et al., 2022; Paul et al., 2023). This is in part understandable since cell cycle phase strongly influences clustering and downstream data analyses (as shown e.g., in our own results; Fig. 1b). Yet it carries a cost when EMT is the object of study: regressing out cell cycle as a technical confound risks attenuating real EMT signal, including the G1-specific genes described here, many of which are absent from canonical EMT lists (Fig. 5b).

EMT–cell cycle coupling has therapeutic implications. Conventional chemotherapy preferentially kills actively dividing cells and may therefore spare the G1 cells with the highest EMT expression, linked to metastatic dissemination. Our model predictions highlight this danger: under simulated cycle-selective chemotherapy, the G1 intermediate state is enriched most consistently (Section 3.6) — the population with the strongest EMT upregulation in our data, and the EMT state most associated with metastatic competence (Pastushenko et al., 2018; Simeonov et al., 2021). Eliminating these cells would require strategies that complement cytotoxicity. TGF-*β* inhibitors offer one such route, reversing EMT phenotype in ovarian cancer cells (Ameri et al., 2019), reducing invasion and metastatic dissemination in pancreatic cancer *in vivo* (Melisi et al., 2008), and decreasing breast cancer metastasis *in vivo* with combinatorial treatment (S.-Y. Park et al., 2015; Vinod et al., 2024).

Several limitations of this study warrant consideration. We study only one cell line (MCF10A) and one EMT-inducing stimulus (TGF-*β*), although EMT is heterogeneous and context-dependent. The results we report also rely on computational inference of cell states rather than direct perturbation. RNA velocity models carry their own assumptions about transcriptional kinetics and treat splicing as a proxy for future state. As with scRNA-seq analysis more generally, an inevitable (often large) number of user-chosen parameters are involved. Single-cell transcriptomics also cannot directly capture certain properties of the cytoskeleton that arise from post-translational regulation (Cho & Kim, 2026). Extending this analysis to additional cell lines, stimuli, and *in vivo* settings would help establish generalizability. Functional cell assays, for example, that test whether cytoskeletal-remodeling inhibitors slow EMT *selectively* in G1 cells could establish causality. The mathematical model considered is highly simplified: extensions could consider more realistic coupling between cell cycle and EMT regulatory networks, at the expense of increased parameter space. The role of extrinsic cell signaling on EMT state represents another future avenue for modeling (Rommelfanger & MacLean, 2021).

Cell cycle phase is not merely a confounding factor to control for in studies of EMT. Rather, it shapes EMT state and dynamics in its own right. We urge for greater consideration of cell cycle effects as a source of biological variation in studies of EMT and beyond.

## Author Statements

### Software and Data Availability

The scRNA-seq datasets analyzed in this study are publicly available from the Gene Expression Omnibus under accession numbers GSE213753 (Panchy et al., 2022) and GSE194019 (Paul et al., 2023). All code used for data analysis and modeling is available on GitHub at: https://github.com/maclean-lab/EMT-cellcycle.

## Acknowledgments

The authors thank members of the MacLean lab for valuable discussions.

## Author Contributions

Conceptualization: MM, AD, RM, ALM. Methodology: MM, AD, ALM. Software: MM, AD, RM. Investigation: MM, AD, RM. Formal analysis: MM, AD, ALM. Visualization: MM, AD. Writing - original draft: MM. Writing - review and editing: MM, AD, ALM. Supervision: ALM. Funding acquisition: ALM.

