## Supplementary Figures for "Cell cycling throttles epithelial–mesenchymal transition"

### 1 SUPPLEMENTARY FIGURES

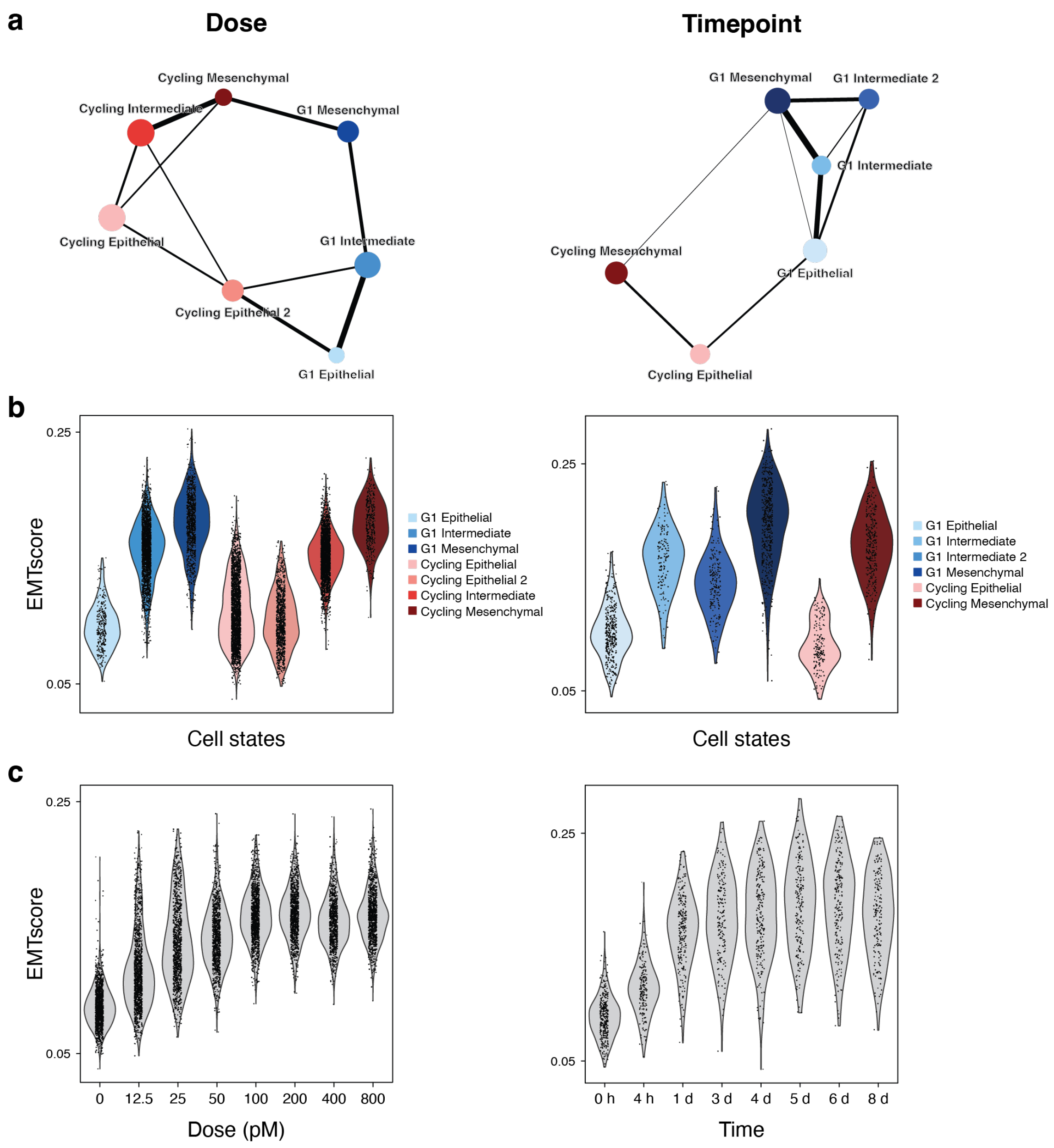


**Supp. Figure 1:** *Cell state structure and EMT progression in the dose and timepoint datasets.* ***a)*** *PAGA graph (Wolf et al., 2019) of cell state connectivity. G1 states are highly connected to one another, as are cycling states; however, G1 intermediate states share no connections with cycling states, indicating transcriptional dissimilarity.* ***b)*** *EMT score by cell state. G1 intermediate and G1 mesenchymal states score higher than their cycling counterparts.* ***c)*** *EMT score by TGF-$\beta$ dose or treatment duration. EMT score increases with both.*

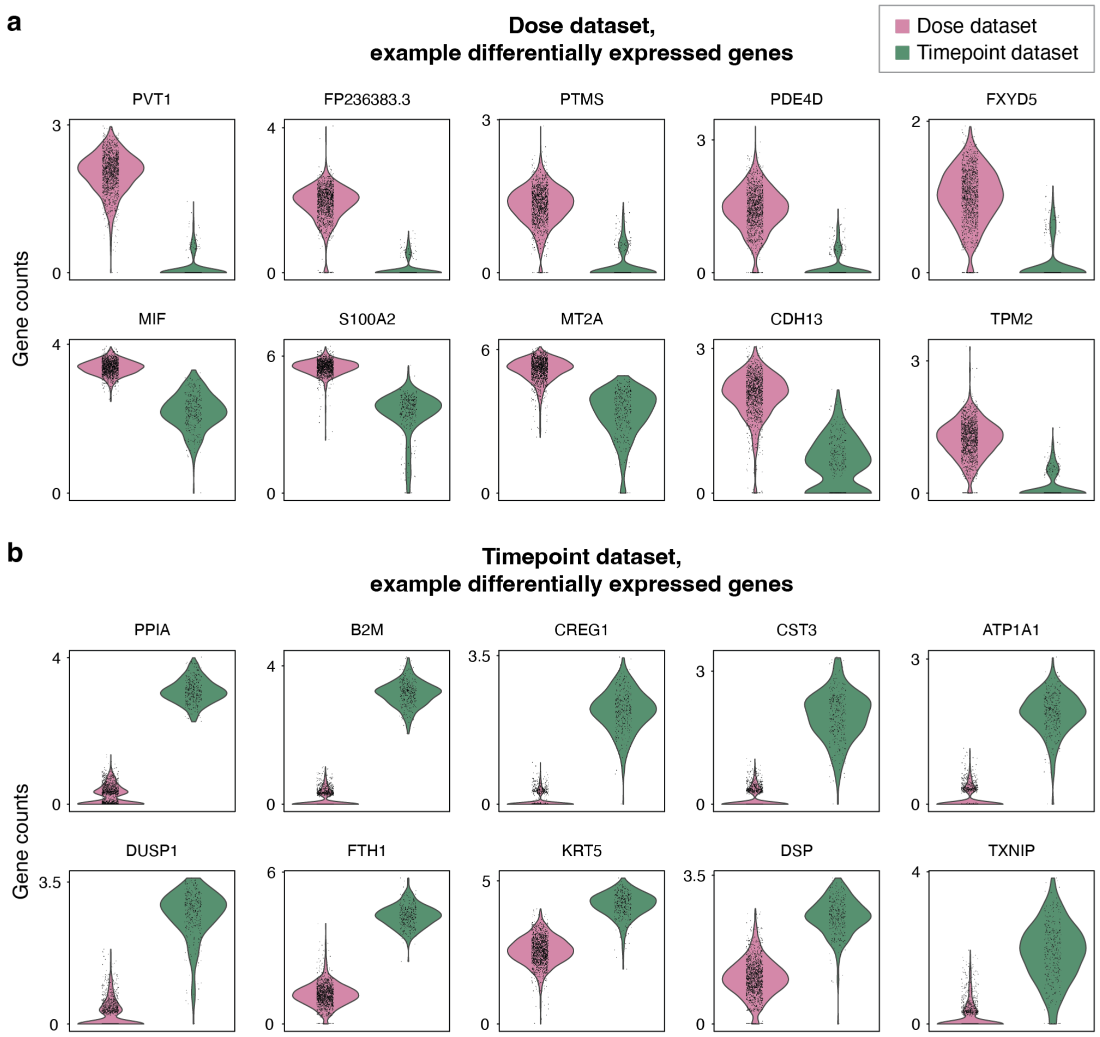


**Supp. Figure 2:** *Transcriptional differences between untreated MCF10A cells in the* ***a)*** *dose and* ***b)*** *timepoint datasets. Expression is shown after total-count normalization, batch integration, and restriction to genes shared between datasets. Many genes are well captured, indicating that the differences are not attributable to dropout alone. Genes shown are examples from several hundred DE genes. Although the datasets differ in cell cycle composition (Fig. 1a), none of the shown genes are cell cycle-associated, suggesting that baseline differences extend beyond cell cycle.*

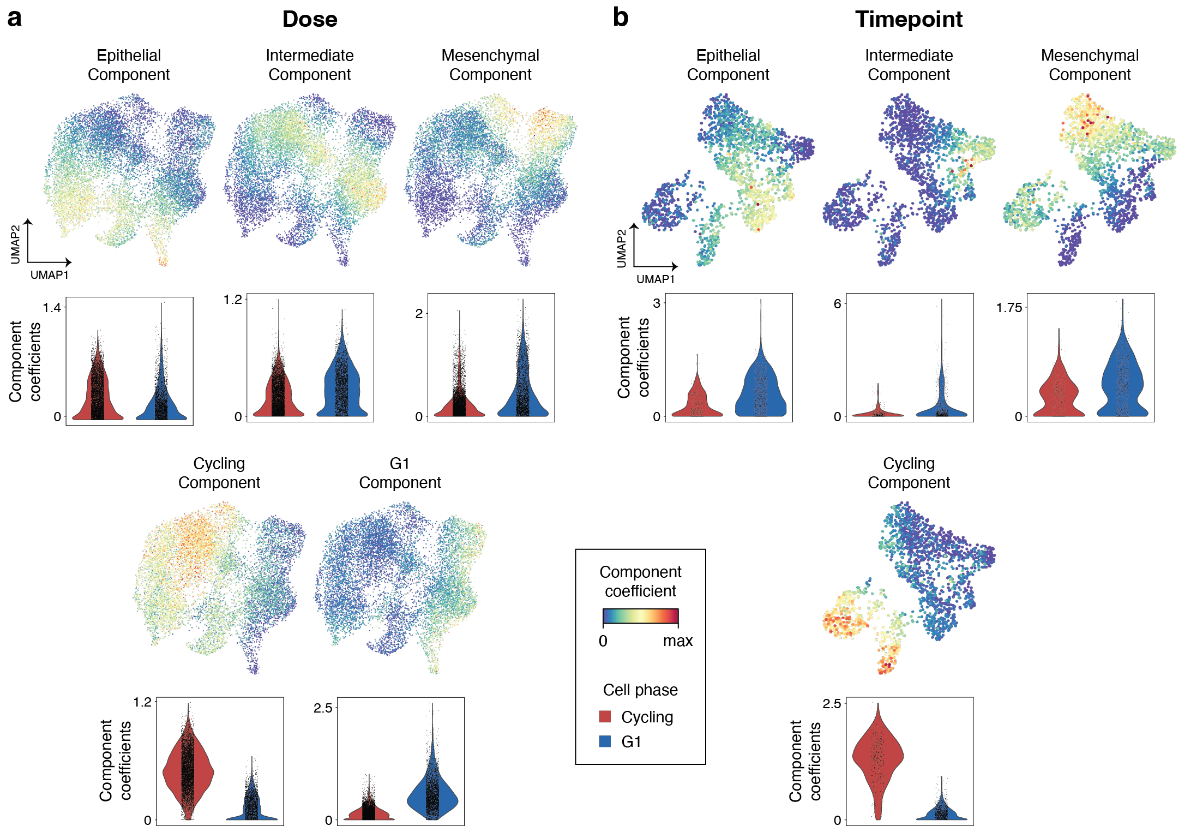


**Supp. Figure 3:** *Non-negative matrix factorization (NMF) of the* ***a)*** *dose and* ***b)*** *timepoint datasets. Both datasets yield epithelial, intermediate, and mesenchymal components; the dose dataset additionally has G1 and cycling components, and the timepoint dataset has a cycling component. Violin plots show per-cell component coefficients in G1 versus cycling cells. Intermediate and mesenchymal coefficients are elevated in G1 cells in both datasets. The number of components was incremented until an additional component no longer corresponded to a distinct cell state.*

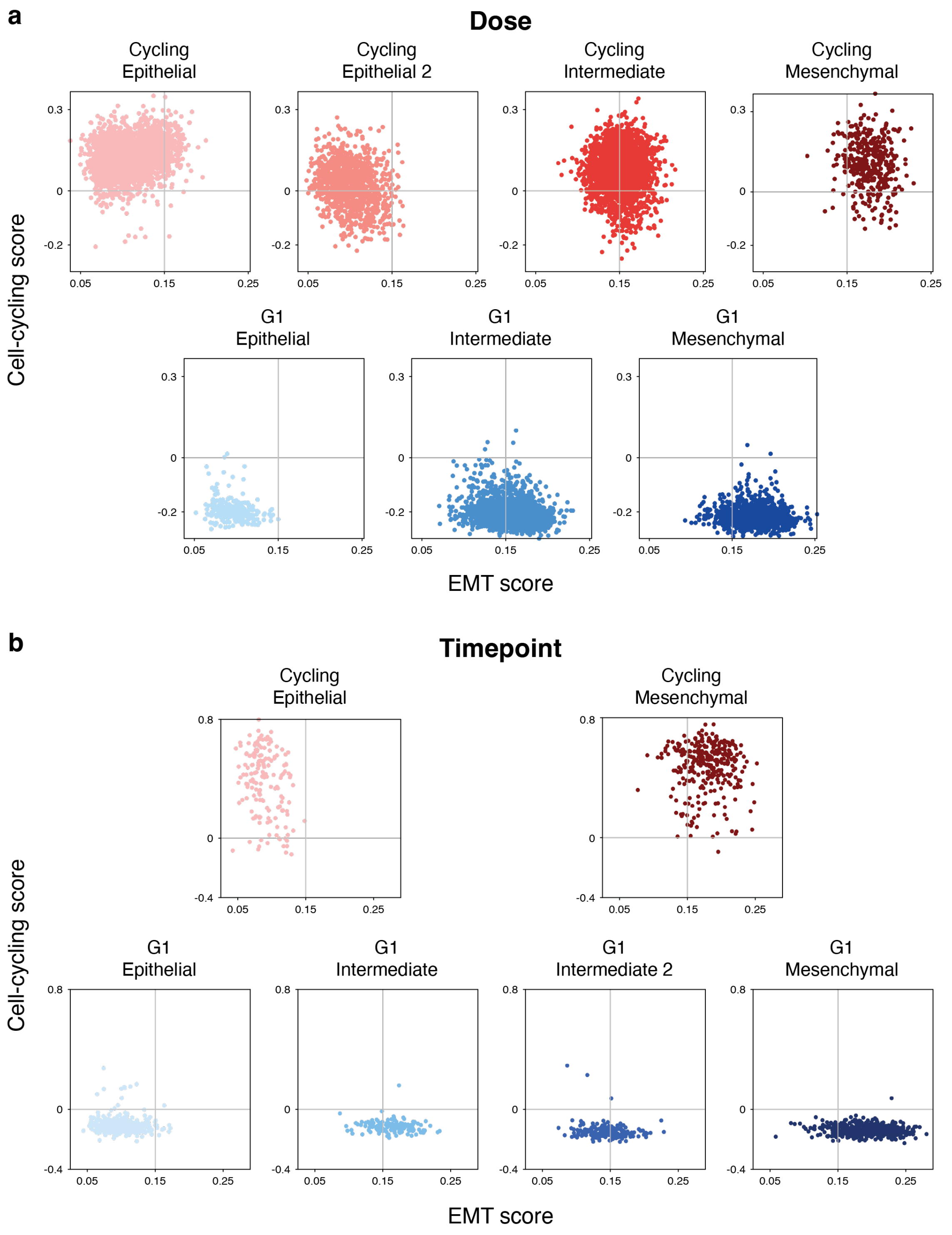


**Supp. Figure 4:** *EMT score and cell-cycling score of single cells, separated by cell state, in the **a)** dose and **b)** timepoint datasets. G1 states have lower cell-cycling scores than cycling states, and EMT score increases from epithelial to intermediate to mesenchymal states.*

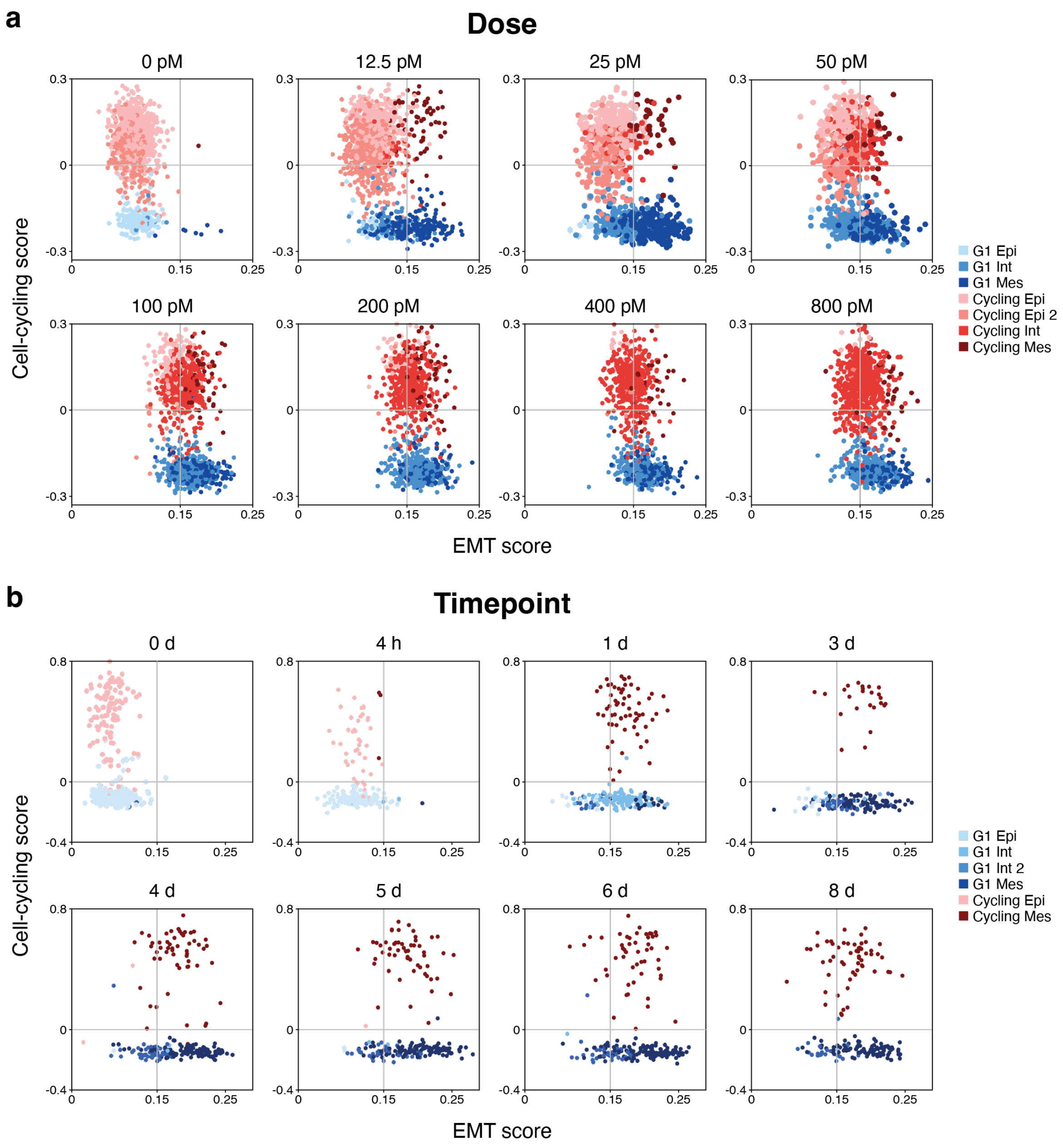


**Supp. Figure 5:** *EMT score and cell-cycling score of single cells by* ***a)*** *TGF-$\beta$ dose and* ***b)*** *treatment duration. EMT score increases with both dose and duration; G1 and cycling cells remain separated along the cell-cycling axis in all conditions.*

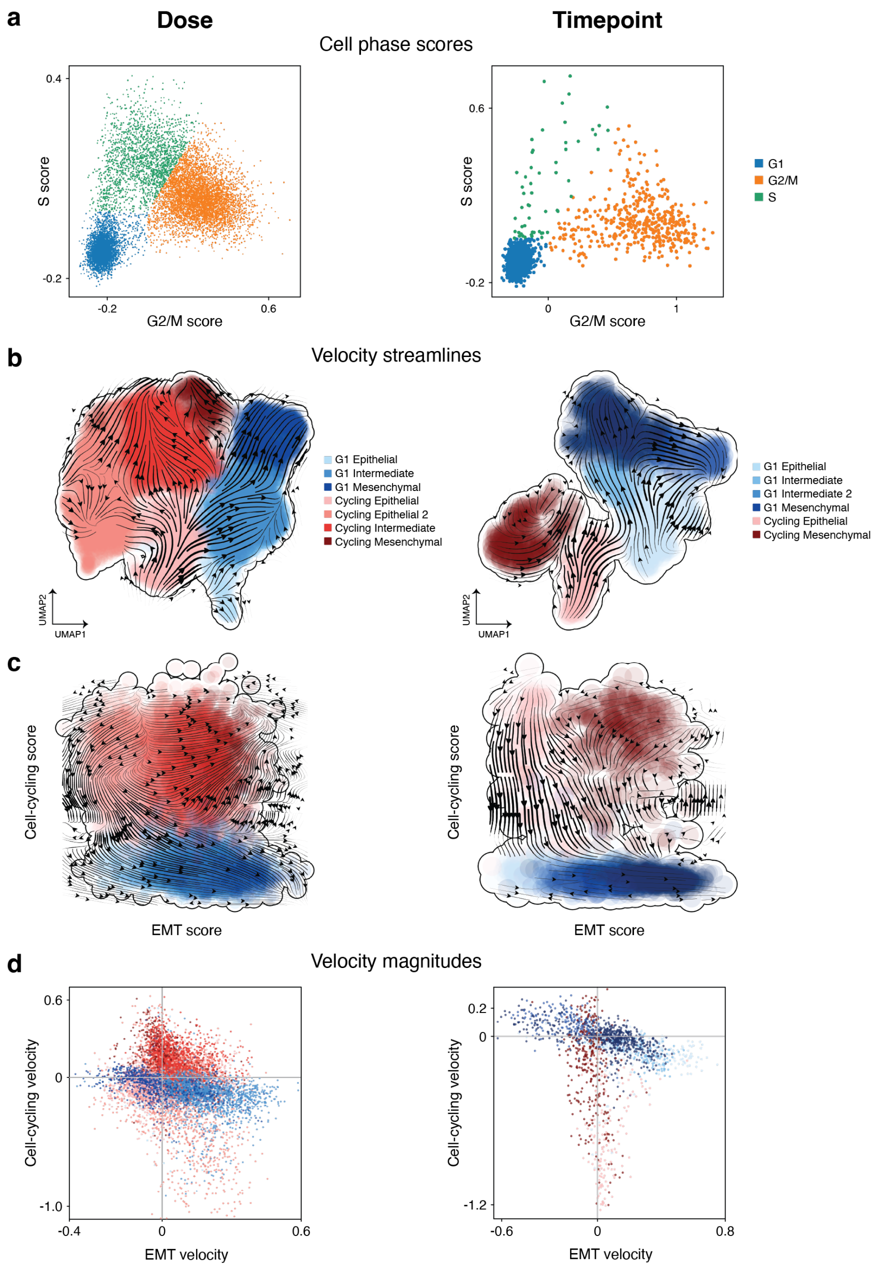


**Supp. Figure 6:** *Cell cycle scores and RNA velocity in the dose and timepoint datasets.* ***a)*** *S and G2/M scores, computed with Scanpy's* `score_genes_cell_cycle` *using the gene sets of Tirosh et al. (2016). G1 cells group together, whereas S and G2/M cells form a continuum.* ***b)*** *Velocity streamlines on UMAP.* ***c)*** *Velocity streamlines in EMT–cell-cycling score space (see Methods).* ***d)*** *Per-cell velocity magnitudes in EMT–cell-cycling score space, colored by phase (G1, blue; cycling, red). G1 and cycling cells show distinct velocity distributions, with G1 cells reaching higher positive EMT velocities.*

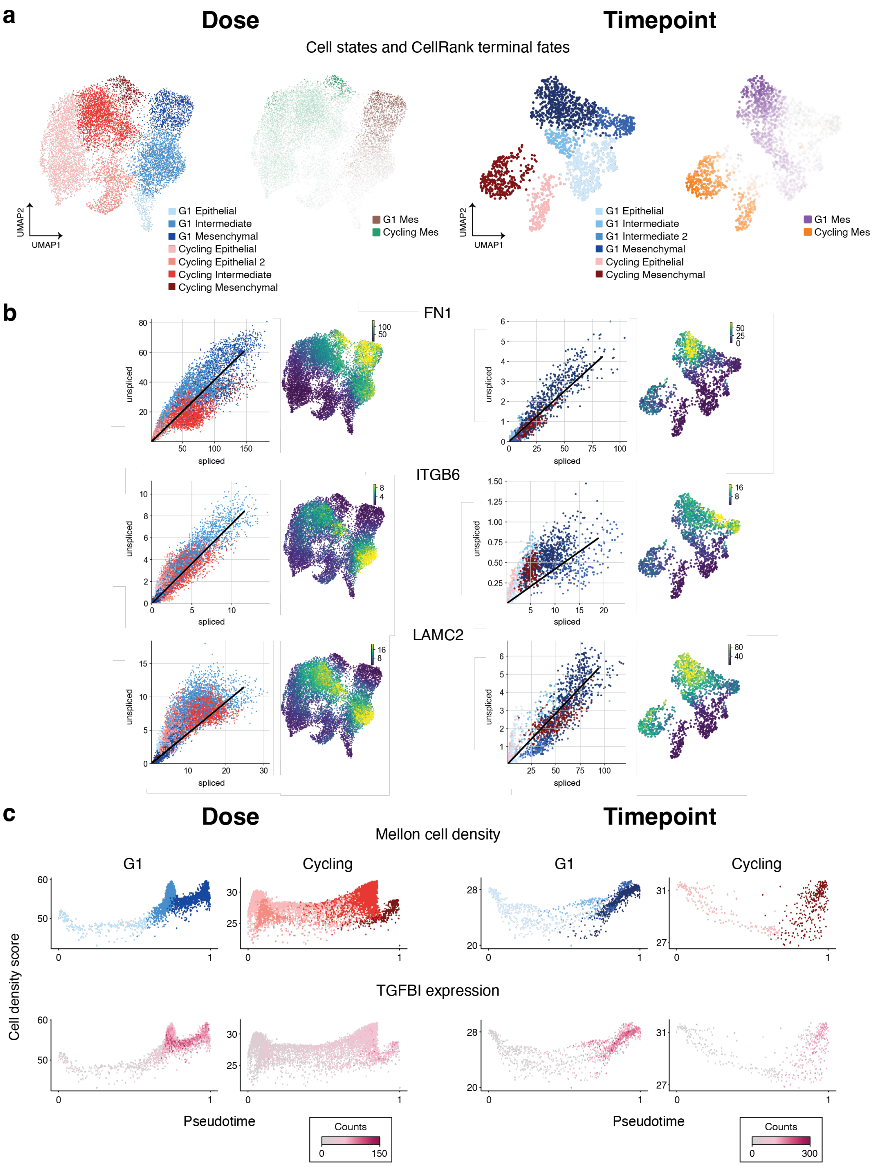


**Supp. Figure 7:** *Identification of G1 EMT-associated genes using RNA velocity and cell-state density.* ***a)*** *CellRank fate probabilities toward the G1 mesenchymal and cycling mesenchymal terminal states, which were identified from a combined RNA velocity and expression kernel. Genes were ranked by their correlation with these fate probabilities.* ***b)*** *Examples of velocity genes (FN1, ITGB6, LAMC2) upregulated in G1 EMT, using CellRank and scVelo.* ***c)*** *Cell-state densities computed with Mellon. In the dose dataset, epithelial, intermediate, and mesenchymal states each form a distinct density peak. In the timepoint dataset, epithelial and mesenchymal states also form peaks; G1 Intermediate 2 forms a high-density peak, suggesting a stable state, whereas G1 Intermediate has lower density, suggesting a more transient one. Overall, transition genes were identified as those whose expression changes most in low-density regions. TGFBI is shown as an example in both datasets.*

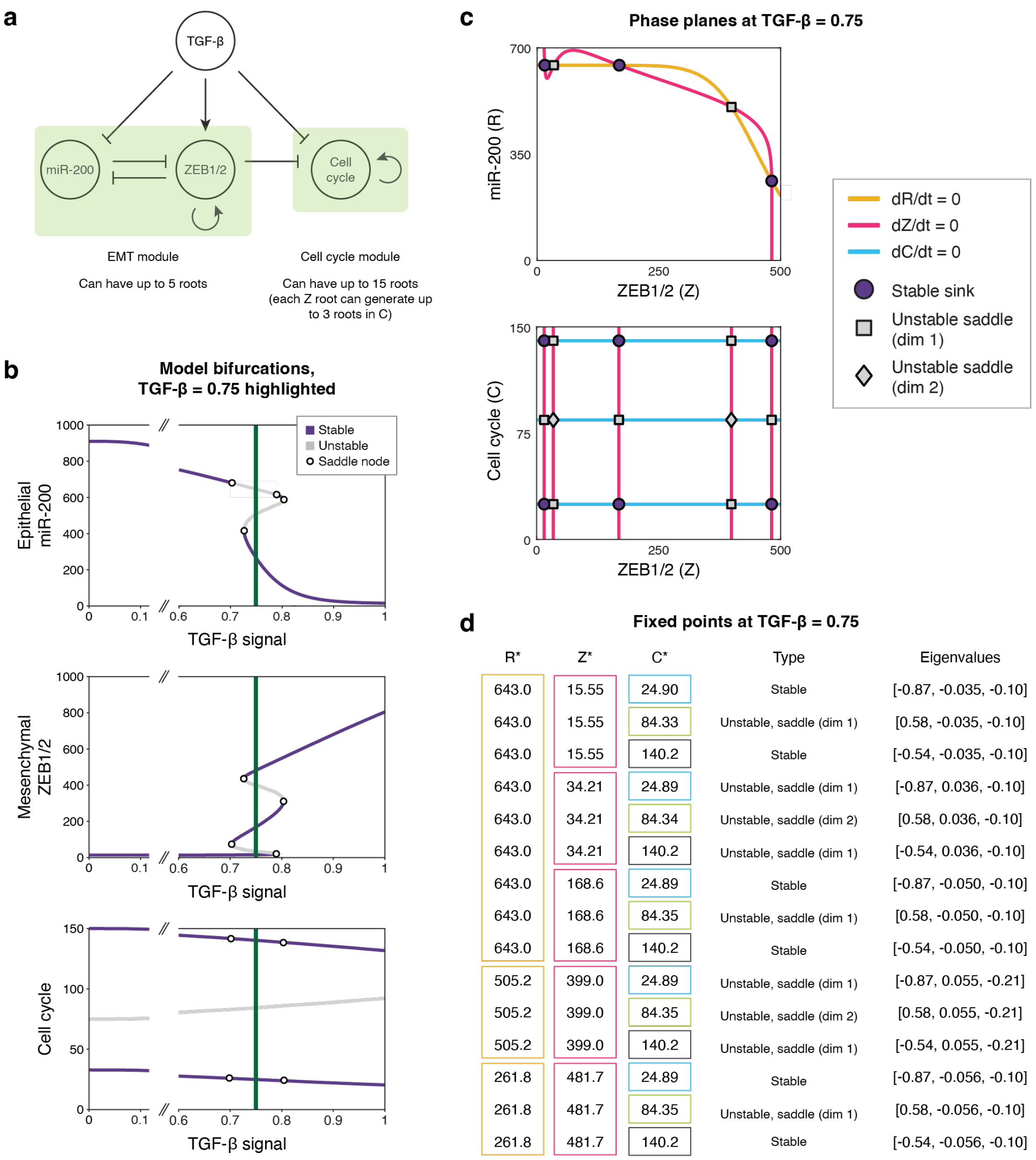


| R* | Z* | C* | Type | Eigenvalues |
|---|---|---|---|---|
| 643.0 | 15.55 | 24.90 | Stable | [-0.87, -0.035, -0.10] |
| 643.0 | 15.55 | 84.33 | Unstable, saddle (dim 1) | [0.58, -0.035, -0.10] |
| 643.0 | 15.55 | 140.2 | Stable | [-0.54, -0.035, -0.10] |
| 643.0 | 34.21 | 24.89 | Unstable, saddle (dim 1) | [-0.87, 0.036, -0.10] |
| 643.0 | 34.21 | 84.34 | Unstable, saddle (dim 2) | [0.58, 0.036, -0.10] |
| 643.0 | 34.21 | 140.2 | Unstable, saddle (dim 1) | [-0.54, 0.036, -0.10] |
| 643.0 | 168.6 | 24.89 | Stable | [-0.87, -0.050, -0.10] |
| 643.0 | 168.6 | 84.35 | Unstable, saddle (dim 1) | [0.58, -0.050, -0.10] |
| 643.0 | 168.6 | 140.2 | Stable | [-0.54, -0.050, -0.10] |
| 505.2 | 399.0 | 24.89 | Unstable, saddle (dim 1) | [-0.87, 0.055, -0.21] |
| 505.2 | 399.0 | 84.35 | Unstable, saddle (dim 2) | [0.58, 0.055, -0.21] |
| 505.2 | 399.0 | 140.2 | Unstable, saddle (dim 1) | [-0.54, 0.055, -0.21] |
| 261.8 | 481.7 | 24.89 | Stable | [-0.87, -0.056, -0.10] |
| 261.8 | 481.7 | 84.35 | Unstable, saddle (dim 1) | [0.58, -0.056, -0.10] |
| 261.8 | 481.7 | 140.2 | Stable | [-0.54, -0.056, -0.10] |

**Supp. Figure 8:** *Model overview with phase planes and fixed points (FPs).* ***a)*** *Model diagram with the EMT and cell cycle subsystems highlighted.* ***b)*** *Bifurcation diagrams (as in Fig. 6b) with FPs at TGF-$\beta = 0.75$ highlighted.* ***c)*** *Example phase plane at TGF-$\beta = 0.75$. FP values vary with TGF-$\beta$, but the phase plane is qualitatively similar across the tristable range (TGF-$\beta \approx 0.7$–$0.8$), which has the maximal number of FPs. Although there are 15 FPs, their values overlap when projected onto a single variable: miR-200 has 3 distinct FP values, ZEB1/2 has 5, and the cell cycle module has 3. The bifurcation diagrams in (b) therefore show at most 3 or 5 branches.* ***d)*** *Values of all 15 FPs at TGF-$\beta = 0.75$; boxes indicate value overlap.*

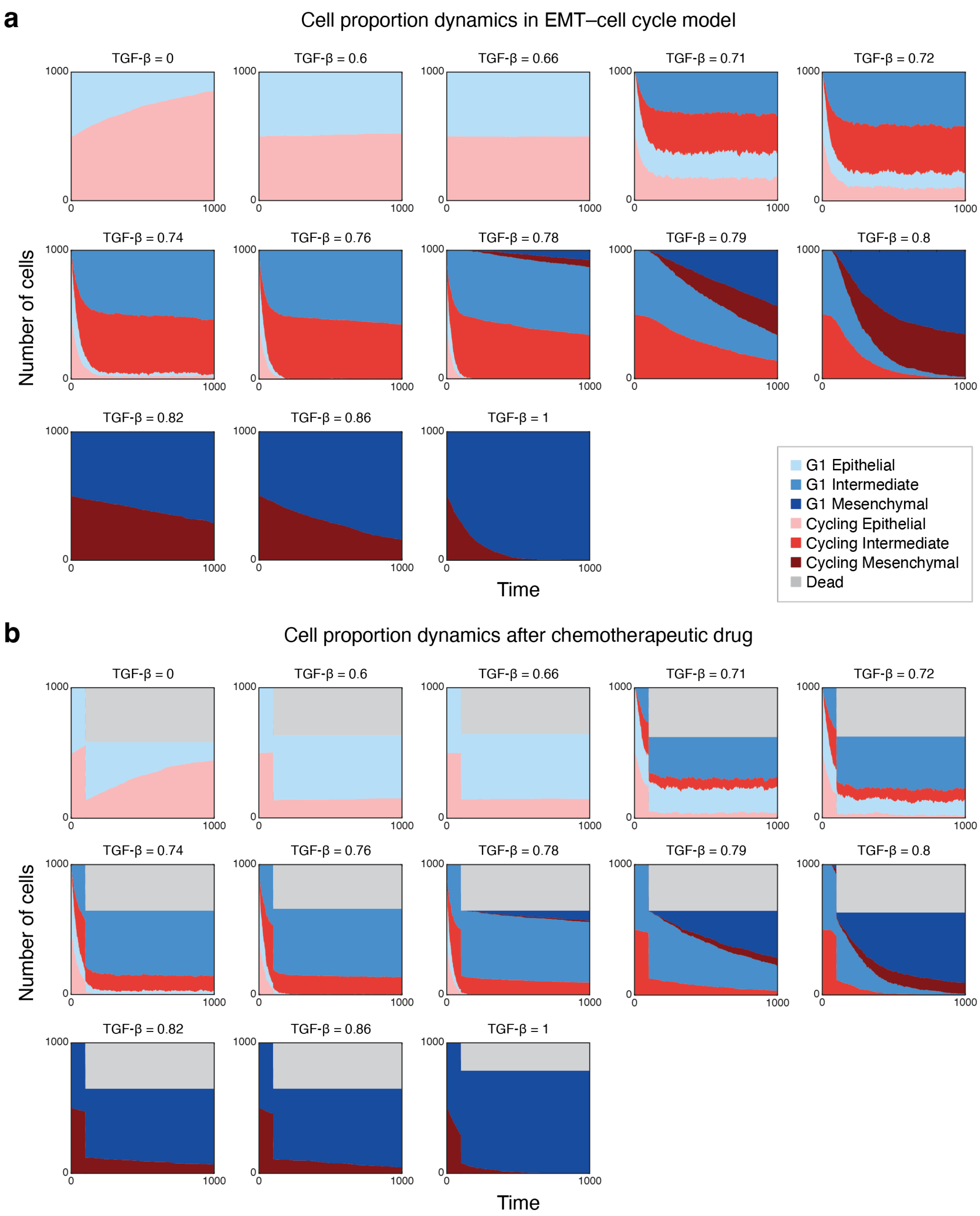


**Supp. Figure 9:** *Population trajectories of the model simulations (steady states shown in Fig. 6c,d). Each subplot corresponds to a single TGF-$\beta$ level.* ***a)*** *Untreated.* ***b)*** *Modeled chemotherapeutic response, in which 75% of cycling cells are removed at $t = 100$. At TGF-$\beta = 0$, treated populations recover to a distribution similar to that of untreated populations. At higher TGF-$\beta$ levels, treated populations shift toward G1 states, especially G1 intermediate. G1 mesenchymal cells also increase, although their relative gain (Fig. 6e) is smaller at the highest TGF-$\beta$ levels, where G1 cells already predominate before treatment.*
